# Locus coeruleus neuromelanin accumulation and dissipation across the lifespan

**DOI:** 10.1101/2023.10.17.562814

**Authors:** Elizabeth Riley, Nicholas Cicero, Khena Swallow, Eve De Rosa, Adam Anderson

**Affiliations:** Department of Psychology, Cornell University

**Keywords:** neuromelanin, locus coeruleus, aging, cognition

## Abstract

The pigment neuromelanin, produced in the locus coeruleus (LC) as a byproduct of catecholamine synthesis, gives the “blue spot” its name, and both identifies LC neurons and is thought to play an important yet complex role in normal and pathological aging. Using neuromelanin-sensitive T1-weighted turbo spin echo MRI scans we characterized volume and neuromelanin signal intensity in the LC of 96 participants between the ages of 19 and 86. Although LC volume did not change significantly throughout the lifespan, LC neuromelanin signal intensity increased from early adulthood, peaked around age 60 and precipitously declined thereafter. Neuromelanin intensity was greater in the caudal relative to rostral extent and in women relative to men. With regard to function, rostral LC neuromelanin intensity was associated with fluid cognition in older adults (60+) only in those above the 50th percentile of cognitive ability for age. The gradual accumulation of LC neuromelanin across the lifespan, its sudden dissipation in later life, and relation to preserved cognitive function, is consistent with its complex role in normal and pathological aging.

## Introduction

The locus coeruleus, located in the pons, is part of the reticular activating system and the major source of modulatory neurochemical norepinephrine for the brain (Aston-Jones & Cohen, 2005; Berridge & Waterhouse, 2003; Sara & Bouret, 2012). Research has linked the status of the locus coeruleus norepinephrine system to cognitive health and the underpinnings of multiple neurodegenerative diseases (Feinstein et al., 2016), including Alzheimer’s Disease (Beardmore et al., 2021; Cassidy et al., 2022; Chan-Palay & Asan, 1989) and Parkinson’s Disease (Gesi et al., 2000; Zarow et al., 2003). The LC is vulnerable to pathological change extremely early in the disease process (Braak et al., 2011; Cassidy et al., 2022), especially in its accumulation of hyperphosphorylated tau (Chalermpalanupap et al., 2017), which can spread throughout its extensive dendritic arbor to the cortex (Ghosh et al., 2019). In addition to this role in spreading pathology, the loss of LC neurons results in dysfunction due to the crucial role that the norepinephrine system plays in sleep (Van Egroo et al., 2022), arousal (Ross & Van Bockstaele, 2021), attention (Plini et al., 2021), cognitive reserve (Clewett et al., 2015), and cognitive control (Aston-Jones et al., 2000). Pathological changes in the LC take place years or even decades before symptomatic illness and may be detectable in those at risk for disease (Granholm et al., 2017; Kremen et al., 2019; Plini et al., 2021). For all these reasons, there is great interest in assessing the LC in vivo so that disease states may be predicted or prevented.

LC measurement in vivo rests primarily on neuromelanin-sensitive MRI sequences (Clewett et al., 2015; Sulzer et al., 2018; Trujillo et al., 2023). Neuromelanin is a pigment that is produced as a byproduct of catecholamine synthesis, in a reaction that requires iron and results in the storage of iron-containing neuromelanin polymers in permanent granules within neurons (Nelson et al., 2009; Shibata et al., 2006; van der Pluijm et al., 2021). Neuromelanin cannot be efficiently cleared by the cell, and thus builds up throughout the lifespan, at least until neurons are lost (Haining & Achat-Mendes, 2017). LC can be reliably visualized as hyperintense on MRI with neuromelanin-specific sequences (Langley et al., 2016).

The relationship between neuromelanin and neuron health is complex. Since neuromelanin serves as a de facto record of former norepinephrine production, it could be said to cumulatively index prior mental engagement and arousal levels (Clewett et al., 2015). Altered LC activity and norepinephrine release are also associated with psychiatric conditions such as depression (Guinea-Izquierdo et al., 2021; Klimek et al., 1997) and anxiety or PTSD (Celebi, 2022; Ross & Van Bockstaele, 2021) In the context of cognitive aging, sustained mental engagement is associated with better cognitive outcomes (Ryff et al., 2016). Likely for this reason, some research has uncovered a positive relationship between neuromelanin intensity and cognitive performance (Clewett et al., 2015), including future cognitive performance (Dahl et al., 2023), in older adults. However, neuromelanin’s complex role within LC neurons complicates this picture.

Granules of neuromelanin can absorb oxidative radicals in the brain, thus protecting the LC from various toxins (Haining & Achat-Mendes, 2017; Moreno-García et al., 2021; Zucca et al., 2017). The LC has inevitable toxin exposure due to its role in making dopamine and norepinephrine, both of which progress to highly toxic metabolites (Goldstein et al., 2013; Kang et al., 2020), despite their necessity and benefits to cognition. Additionally, the LC accumulates environmental toxins such as heavy metals more than surrounding structures, potentially due to increased exposure through its extensive dendritic arbor in addition to its ability to sequester such toxins (Pamphlett et al., 2018, 2020). These toxins remain sequestered in the neuromelanin granules indefinitely (Haining & Achat-Mendes, 2017; Iannitelli et al., 2023). If the granules are damaged, toxins then spill out resulting in inflammation and cell death (Zecca et al., 2003; Zucca et al., 2017). The conditions that cause damage to LC neuromelanin granules appear to include hyperphosphorylated tau accumulation (Jacobs et al., 2019), and at least in animal models, and an overaccumulation of neuromelanin itself, beyond some unknown saturation point (Carballo-Carbajal et al., 2019; Iannitelli et al., 2023). Thus, the presence of neuromelanin granules is neither unequivocally good or bad, and their value or risk to LC neurons is difficult to determine especially in older adults.

Within an individual, there may come a tipping point, before which neuromelanin accumulation is the happy result of cumulative mental engagement which provides detoxification of a vital brain nucleus, and after which neurons are being lost due to overload of either toxins or neuromelanin itself. This may explain why some previous post mortem and in vivo studies have detected a peak in LC neuromelanin around age 50-60 (Liu et al., 2019; Manaye et al., 1995; Mann et al., 1985; Shibata et al., 2006). A similar inverted-U life course has been found for the substantia nigra (Xing et al., 2018), the other neuromelanin-containing structure in the brain. For older adults, larger amounts of remaining neuromelanin may not necessarily be beneficial, but may indicate that such individuals have not yet reached their neuromelanin tipping point, leaving their LC vulnerable.

In attempting to understand the diagnostic or prognostic value of neuromelanin imaging, it is crucial to understand the typical life course of this enigmatic molecule, and to attempt to refine the relationship between its presence and cognitive function. Recent efforts by others have strongly suggested an inverted-U shaped curve of neuromelanin intensity across the lifespan (Liu et al., 2019). This result is important to replicate particularly from middle to older age. Given demographic disparities in Alzheimer’s related dementias it is also imperative to examine groups from diverse racial backgrounds (Babulal et al., 2019). Most complex is understanding the relationship between neuromelanin intensity and cognition, which is unclear, given neuromelanin can represent both a protective factor and a vulnerability depending on the life stage. These important factors deserve exploration. To that end, we present an analysis of the LC that maximizes neuromelanin contrast to characterize its LC volume and neuromelanin signal intensity in 96 individuals aged 19-86, with comparisons by sex and race, and differences by cognitive functioning, using a method of neuromelanin intensity measurement (Turker et al., 2021) with high sensitivity and reliability across the lifespan.

## Results

### Localization of the LC

We identified the locus coeruleus (LC) in 96 participants between the ages of 19-86 using neuromelanin-sensitive turbo spin echo T1-weighted MRI scans. The underlying mechanism for contrast on such scans remains unclear (Oshima et al., 2021; Priovoulos et al., 2020) but has previously thought to be related to the T1-shortening effects of neuromelanin, thanks to its iron binding ability (Sulzer et al., 2018). All LC intensity and volume measurements were made using individually hand-drawn ROIs made by two independent raters for each participant according to our previously published methods (Turker et al., 2021). Using the individual LC ROIs, we created a probabilistic map of the location of the locus coeruleus (LC) in MNI space. The probabilistic map spanned from coronal coordinates y = −39 to −32, axial coordinates z = −32 to −21, and sagittal coordinates x = −7 to −2 and x = 3 to 8 (MNI coordinates, **Figure 1**).

**Fig. 1.**
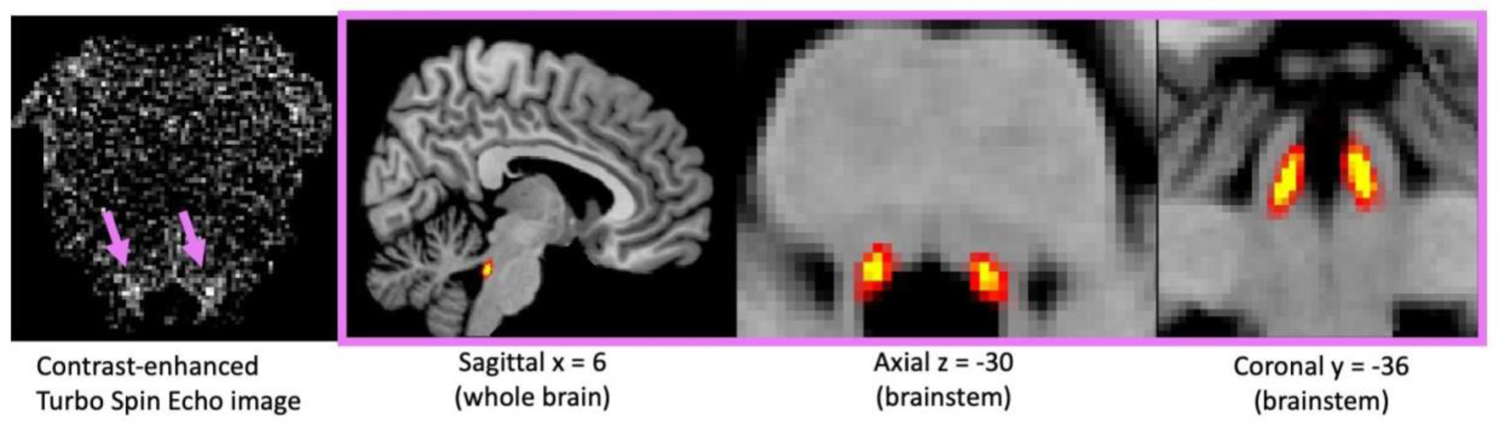
On the left, an example of contrast-enhanced turbo spin echo image in (axial view, limited to brainstem) for localization of the locus coeruleus (bright spots indicated by purple arrows). On the right, three views of a probabilistic map of LC location using data from 96 individuals. MNI coordinates.

### LC volume

The average total LC volume (both left and right) across all participants was 37.3 mm3, with a SD of 22.2 mm3 and a range from 6.1 to 133.1 mm3 (**Figure. 2A and 2B**). A linear model (using lme4 in R, REF) tested the effect of age group on total volume. LC volume did not differ significantly between age groups (F(4) = 0.21, p = 0.92).

**Fig. 2.**
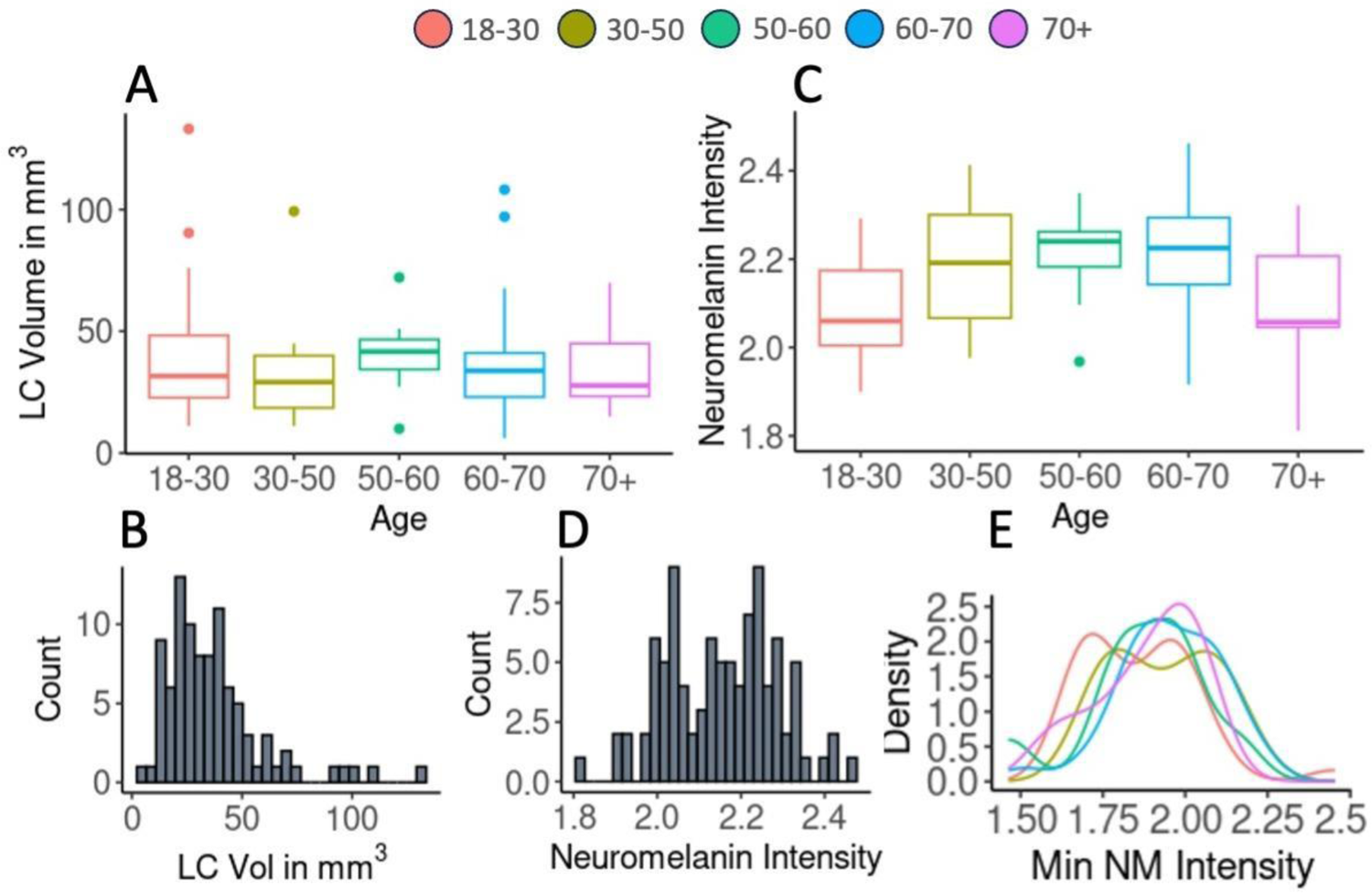
(A) Boxplots showing distribution of total LC volume (left and right combined) across age groups. (B) Histogram of total LC volume for all participants. (C) Boxplots showing distribution of neuromelanin intensity (average per person, including left and right) across age groups. (D) Histogram of average neuromelanin intensity for all participants. (E) Density plot showing distribution of minimum neuromelanin (NM) intensity values by age group. Minimum NM intensity is the least intense voxel that was included in each individual’s hand-drawn LC ROI. In box plots, the middle line shows median, the top and bottom of the box show 75th and 25th percentiles, the top and bottom whiskers show 95th and 5th percentiles, and dots are outliers. Neuromelanin intensity is measured in multiples of the maximum non-LC, non-outlier voxel value in the brainstem.

### LC neuromelanin intensity

Neuromelanin intensity was expressed in terms of multiples of the maximum intensity of non-LC voxels in the same brainstem slices where LC was identified. The average LC neuromelanin intensity across all participants was 2.16, or 2.16 times higher than the calculated maximum of non-LC voxels in the image, with a SD of 0.16 and a range from 1.81 to 2.46 (**Fig. 2C and 2D**). Locus coeruleus neuromelanin intensity increased from young adulthood, was highest around age 60, and declined thereafter (**Fig. 2C**). A linear model was constructed to test the effect of age group on average neuromelanin intensity; intensity varied significantly with age group (F(4) = 4.9, p = 0.0011). Age groups 30-50, 50-60, and 60-70 all differed significantly from the reference age group 18-30 (all t > 2.3, all p < 0.03), while age group 70+ did not (t = 0.55, p = 0.57). Since neuromelanin intensity across the lifespan appeared to follow an inverted-U shaped curve, we also fitted a generalized additive model (Sørensen et al., 2021) to test the relationship between age (by year, rather than by group) and neuromelanin intensity, and found a significant relationship (for the age term, edf = 2.49, F = 4.46, p = 0.0053, R squared = 0.13).

To ensure that there was no systematic bias in the measurement of LC neuromelanin intensity across age groups, we tested whether minimum neuromelanin intensity (the lowest-intensity voxel included within each individual’s hand-drawn LC ROI) varied with age. Overall, it did not (F(4) = 1.92, p = 0.12), although the 60-70 age group did have marginally greater minimum voxel values compared with the youngest age group (t = 1.99, p = 0.05). The 70+ age group, in which we saw a marked decrease in average neuromelanin intensity, did not have significantly decreased minimum neuromelanin intensity. The distribution of minimum voxel values was similar in all age groups and did not match the distribution of maximum voxel values (**Fig. 2E**). Furthermore, our Dice similarity coefficients for the creation if individual LC ROIs did not differ between younger (< 60) and older (>60) participants (see Methods).These reassuring analyses suggested that our measurements of volume were not affected by systematically including less-intense voxels in some groups vs. others.

### Differences between left and right LC

In all age groups, the average volume of the left LC was observed to be larger than the volume of the right LC. The average volume of the left LC was 19.8 mm3, while the average volume of the right LC was 16.6 mm3. A linear model was constructed to test whether side affected volume in any age groups; the effect of side on volume was marginally significant (F(1) = 3.15, p = 0.07, Fig. 3A and 3B), but there was no interaction between age group and side on volume (F(4) = 0.26, p = 0.89). A similar model was constructed to test whether the left and right LC intensity differed significantly in any age group; they did not (F(1) = 0.14, p = 0.69, Figs. 3C and 3D).

**Fig. 3.**
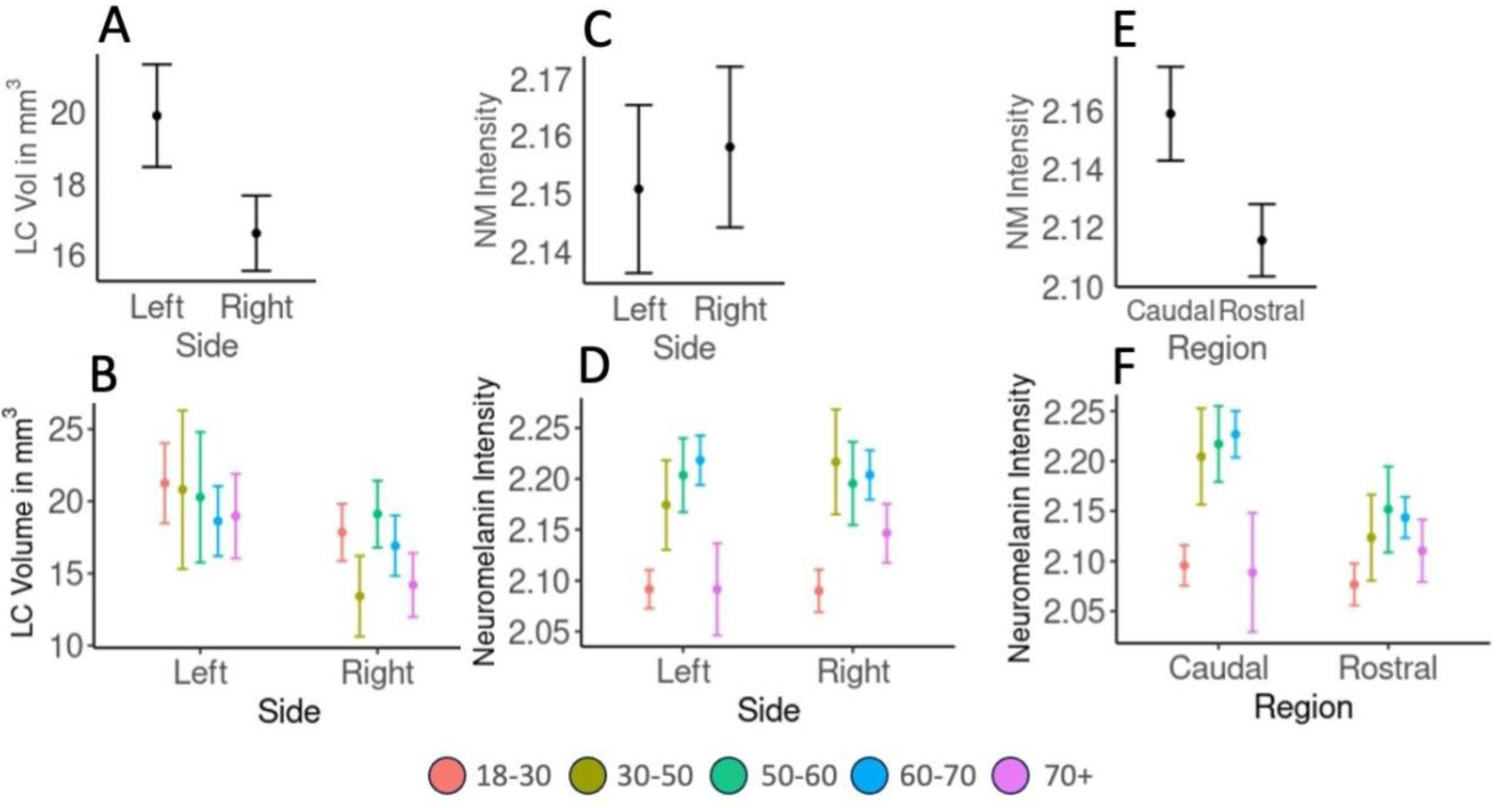
At left, comparison of volume of left and right LC, overall (A) and by age group (B). In the middle, comparison of neuromelanin intensity of left and right LC, overall (C) and by age group (D). On the right, comparison of neuromelanin intensity between rostral and caudal LC, overall (E) and by age group (F). All plots show mean and standard error. Neuromelanin intensity is measured in multiples of the maximum non-LC, non-outlier voxel value in the brainstem.

### Rostral vs. caudal LC

Because the intensity of the rostral LC has often been associated with cognition more than the caudal LC (Chen et al., 2023; Dahl et al., 2019), we repeated the analysis of neuromelanin intensity across age groups using rostral LC intensity only, which also varied significantly with age (F(4) = 4.55, p = 0.0021, with age groups 30-50, 50-60, and 60-70 differing significantly from age group 18-30, all p < 0.04, as in the previous model). In all age groups but one, the average intensity of the caudal LC was observed to be greater than the intensity of the rostral LC. A linear model was constructed to determine whether intensity depended on LC region (rostral vs caudal) in any age group; caudal intensity was found to be significantly higher (F(1) = 4.92, p = 0.027). Average caudal intensity was 2.16 with SD = 0.18, while average rostral intensity was 2.12 with SD = 0.16 (Figs. 3E and 3F).

### LC neuromelanin intensity and cognitive performance

We measured fluid cognition using the NIH Toolbox (Fluid Cognition score) and crystallized cognition using the National Adult Reading Test (NART). Across our sample, raw fluid cognition scores declined sharply across the lifespan. A linear model with raw (non-age normalized) fluid cognition standard score as the dependent variable and age as the independent variable demonstrated that cognitive ability depended on age (F(1) = 56.3, p < 0.000001, Pearson correlation coefficient = −0.67). A corresponding model tested whether crystallized cognition (raw NART score) depended on age; it did not (F(1) = 2.43, p = 0.12, Pearson correlation coefficient = 0.16). Thus, as expected (Moran, 2013), fluid cognition declined with age but crystallized cognition did not. Given that we previously found that neuromelanin intensity increased with age (until the clear downturn in our oldest age group, aged 70+), we expected to find a significant negative correlation between rostral neuromelanin intensity and raw fluid cognition scores in those under 70, and this was indeed the case (r = −0.35, one sample t-test t(74) = −3.2, p = 0.0019).

In order to control for the very strong effect of age on fluid cognition, we next asked whether age-normalized fluid cognition scores (national percentile for age) were associated with neuromelanin intensity. Previous work has found an association between neuromelanin intensity and measures of fluid (REF) and crystallized cognition (REF) only in older adults. Since our data suggested a peak in LC intensity around age 60, we considered that age the lowest for inclusion in this analysis. First we ensured that our normalized cognitive measure did not depend on age by constructing a linear model with age-normalized fluid cognition score as the dependent variable and age as the independent variable, and we found no association (F(1) = 0.27, p = 0.6). We then constructed a linear model to test whether age-normalized fluid cognition ability depended on rostral LC intensity. We found no significant effect (Pearson correlation coefficient = 0.05, F(1) = 0.13, p = 0.73, Fig. 4A). However, upon closer examination, we noticed a number of very low-performing individuals, and that low- and high-performing older adults seemed to have a different relationship between neuromelanin intensity and cognition.

**Fig. 4.**
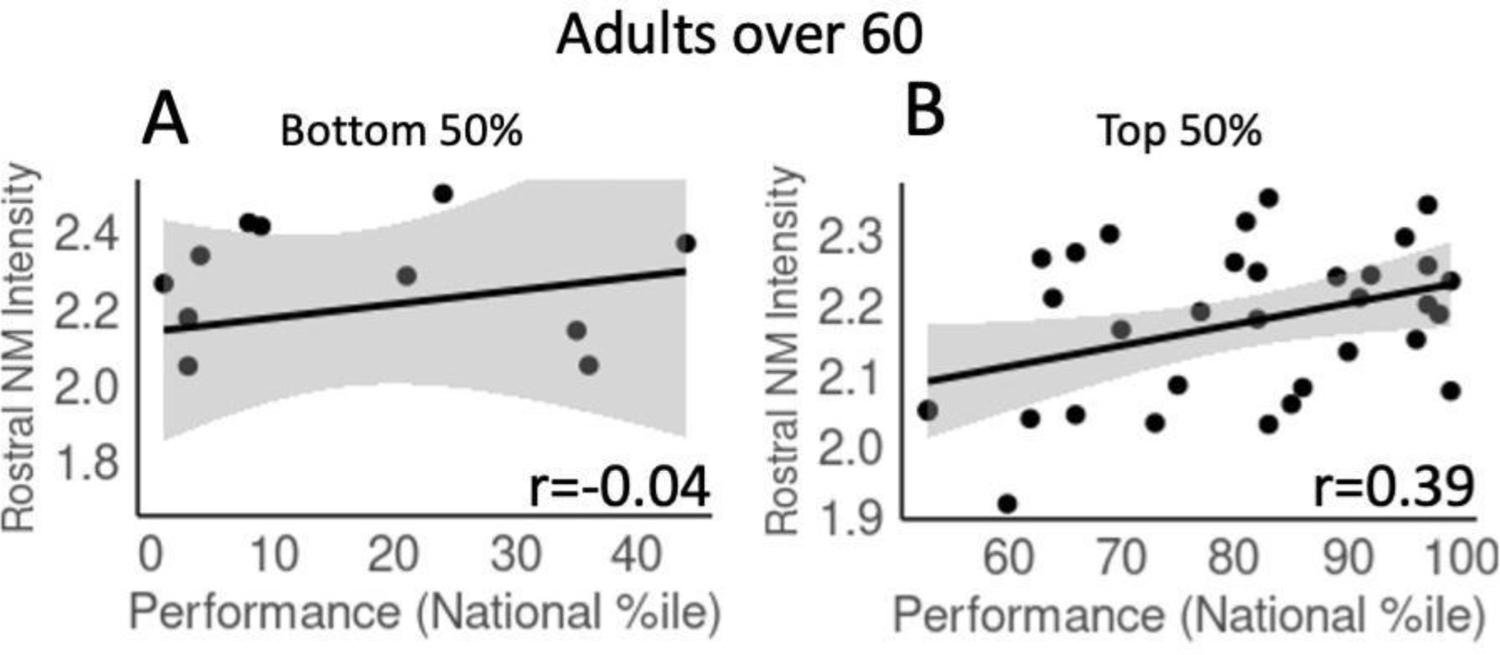
(A) Scatterplot comparing cognitive performance (national percentile for age in fluid cognition composite from NIH toolbox) and rostral LC neuromelanin intensity. (B) Scatterplot comparing cognitive performance amongst above-average (greater than 50th national percentile for age) performers only to rostral LC neuromelanin intensity. (C) Scatterplot comparing cognitive performance amongst below-average (less than 50th national percentile for age) performers only to rostral LC neuromelanin intensity. Lines show linear fit and 95% confidence interval. r is Pearson’s correlation coefficient. Neuromelanin intensity is measured in multiples of the maximum non-LC, non-outlier voxel value in the brainstem.

We repeated the analysis for adults over 60 within each of two cognitive groups (below 50th percentile for age (n=11), and above 50th percentile for age (n=32)). Among adults performing above average for their age, we found a significant positive association between rostral LC intensity and global cognitive performance (Pearson correlation coefficient = 0.39, F(1) = 5.48, p = 0.025, Fig. 4B). However, among adults performing below average for their age, there was no significant relationship (Pearson correlation coefficient = −0.04, F(1) = 0.3, p = 0.57, Fig. 4C). This is despite the fact that high-performing and low-performing groups did not differ in average rostral LC intensity (2.16 ± 0.16 vs 2.16 ± 0.21, respectively).

We used a similar linear model to test whether crystallized cognition (NART score; this score was not corrected for age because it did not depend significantly on age) depended on rostral LC intensity in adults over 60. We found no significant effect in high- or low-performing participants. There was no significant relationship between rostral LC intensity and cognitive performance in adults under 60, nor was there any relationship between caudal LC intensity and cognitive performance in any group. We also found no significant relationship between any cognitive subtests and cognitive performance.

### Neuromelanin intensity differences by race and sex

In all age groups (reduced to 3 because of uneven gender balance within decade age groups), women had higher LC intensity than men. We constructed a linear model to determine whether these differences were significant and whether there was an interaction with age group. There was a significant effect of sex (F(1) = 16.04, p = 0.00012, Fig. 5A) and no significant interaction between sex and age group (F(2) = 0.43, p = 0.65).

**Fig. 5.**
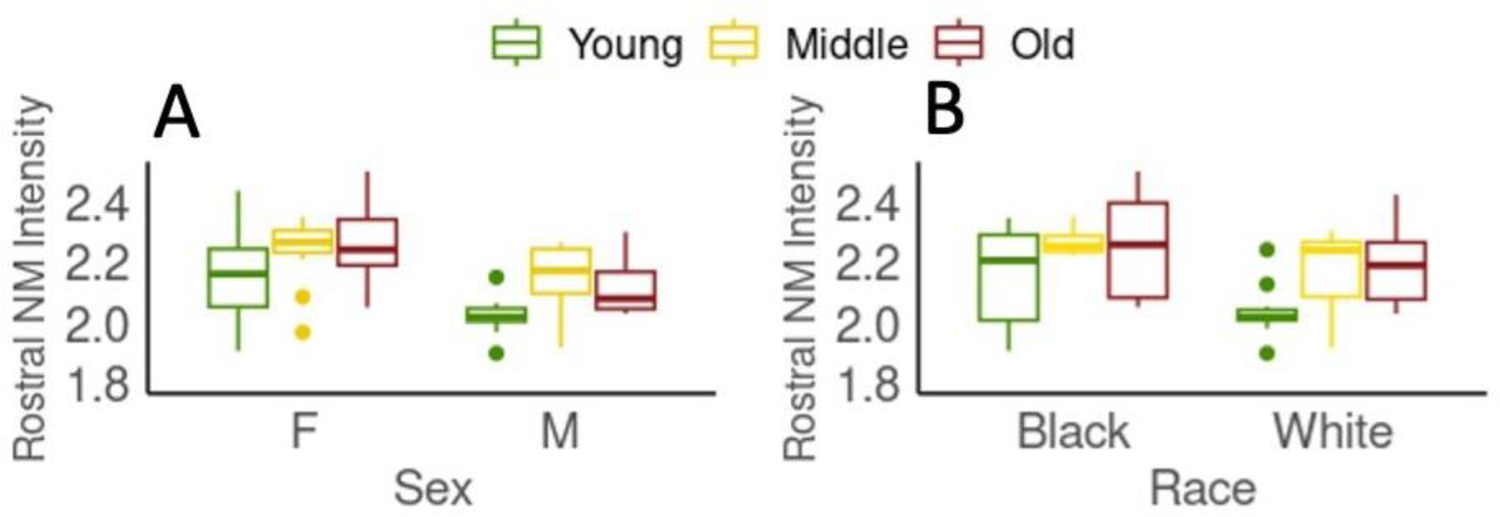
(A) Boxplot showing distribution of rostral LC neuromelanin intensity across condensed age groups and between female and male participants. (B) Boxplot showing distribution of rostral LC neuromelanin intensity across condensed age groups and between Black and White participants. In box plots, the middle line shows median, the top and bottom of the box show 75th and 25th percentiles, the top and bottom whiskers show 95th and 5th percentiles, and dots are outliers. Neuromelanin intensity is measured in multiples of the maximum non-LC, non-outlier voxel value in the brainstem.

We constructed a linear model to test whether there were differences in rostral LC intensity by race, and whether there was any interaction between race and age group. We did not find any significant differences in rostral (or average) LC intensity between participants of black (n= 16) and white (n = 55) race (F(1) = 0.4, p = 0.52, Fig. 5B) in any age group, and there was no significant interaction between race and age group (F(2) = 1.4, p = 0.24). We were unable to test the effect of Asian or Latinx heritage by age group as there were not enough participants in each age category.

## Discussion

Employing neuromelanin-sensitive MRI, we found that neuromelanin intensity, defined relative to the surrounding pons, increased from early adulthood through approximately age 60, and declined thereafter. The caudal portion of the LC had greater neuromelanin intensity than the rostral portion. Demographically, women had greater neuromelanin intensity in the LC than men but we did not find any reliable differences by race. While the caudal portion of the LC had greater intensity, it was the rostral portion of the LC that was associated with cognitive performance. This relation applied only to older adults with above-average cognition for their age and was selective to fluid rather than crystalized measures of cognition.

Previous analyses of neuromelanin across the lifespan have also detected a peak in neuromelanin intensity around age 50-60, including early post mortem studies and two in-vivo MRI studies (Liu et al., 2019; Manaye et al., 1995; Mann et al., 1985; Shibata et al., 2006).

However, other studies have found no change in neuromelanin concentration across the lifespan (Hämmerer et al., 2018; Ohm et al., 1997; Zucca et al., 2006), or simply an increase between younger and older adults (Clewett 2016 in vivo, Betts 2017 in vivo). Thus, even recent examinations of LC neuromelanin intensity have not yet arrived at a completely consistent result. In studies on the substantia nigra, the other neuromelanin-containing brain structure, the inverted-U with a peak around X has been found using neuromelanin-sensitive MRI (Xing et al., 2018). Differences by sex are even less clear, with some analyses (Clewett et al., 2015) showing markedly lower intensity in women, and some showing no difference (Betts et al., 2017; Shibata et al., 2006). By contrast, our analysis shows *increased* signal intensity in women.

Though our sample is relatively small (N = 96), it is larger than the sample size in all studies cited in this paragraph except for (Liu et al., 2019), with which it most closely agrees. That our life course of neuromelanin intensity was internally replicated between left and right LC, between rostral and caudal regions, within racial groups and for both males and females, gives some further confidence in its accuracy. As far as we are aware, no other published studies have examined neuromelanin intensity across different races.

We found a positive association between rostral LC intensity and fluid, but not crystallized, cognition specific to older adults with above-average cognition. Related results have previously been reported by others (Bell et al., 2023; Clewett et al., 2015; Dahl et al., 2019, 2023), though not with perfect alignment. Clewett et al., 2015 found a significant association between crystallized intelligence and overall LC intensity, Bell et al., 2023 found an significant association between subjective cognition and rostral LC intensity, and Dahl et al., 2019 found a significant association between memory performance and rostral LC intensity, and then in 2023 found that LC intensity significantly predicted future episodic memory performance. This growing literature will benefit from continued study with large and diverse sample sizes.

Our study had uneven group sizes, due to the relative difficulty of recruiting middle-aged participants, and while we had significant diversity within our sample, we were unable to ensure that every age, sex, and racial group was matched in all ways that might theoretically impact neuromelanin intensity (e.g. education). Furthermore, even though we detected a significant positive relationship between neuromelanin and fluid cognition, it was not strong or specific enough such that neuromelanin measurement would tell a clear story about an individual.

The cause of loss of neuromelanin in older adults is not clear. It could be either death of catecholaminergic cells, or neuromelanin loss without cell death. It is difficult to distinguish between these two possibilities. There is some evidence for reduced neuromelanin content in surviving cells in older adults (Faucheux et al., 2003; Mann et al., 1985), although neuromelanin loss without cell death is considered less likely because previous work has linked neuromelanin directly to surviving LC neurons (Keren et al., 2009, 2015; Kitao et al., 2013). (This work comes from neuromelanin-sensitive MRI in patients with Parkinson’s disease, a literature which has produced more than a dozen replications of reduced neuromelanin signal, see (Sulzer et al., 2018) Although we measured LC volume, diffuse neuronal loss across the LC could result in an unchanged volume even after substantial cell death. Furthermore, although neuromelanin clearly accumulates in granules within cells, it also exists in the extracellular space, for an undetermined amount of time prior to clearance, likely by microglia (Zhang et al., 2011; Zucca et al., 2006). Recent quantitative susceptibility mapping has made clear that iron resulting from neuromelanin granule loss is taken up by surrounding cells, where it is subsequently visible on MRI using iron-sensitive sequences that are *not* sensitive to neuromelanin (He et al., 2023; Takahashi et al., 2018).

The question of how neuromelanin impacts neuron health, and what to make of its decrease in old age, is being actively investigated and has been labeled a paradox (Moreno-García et al., 2021), an enigma (Youdim et al., 1994) and a riddle (Iannitelli & Weinshenker, 2023). Since neuromelanin unquestionably results from the oxidation of intermediate products in the norepinephrine production pathway, the amount produced might be considered a reporter of total NE and DA production in the locus coeruleus (Clewett et al., 2015), although very recent work suggests the possibility of active neuromelanin clearance (Bachman et al., 2023). Because NE exerts numerous neuroprotective effects on the brain and is essential for cognition (Feinstein et al., 2016; Robertson, 2013), and since neuromelanin is a highly effective quencher of toxins that might otherwise damage the LC (Moreno-García et al., 2021), greater production or preservation of this neurochemical could account for the positive relationship between neuromelanin intensity and cognition, which we also observed. On the other hand, it is very clear from recent animal studies that overexpression of human tyrosinase in both the substantia nigra and LC in animal models results in abundant neuromelanin production and very quickly induces neurodegeneration and associated phenotypes as a direct consequence of neuromelanin accumulation (Carballo-Carbajal et al., 2019; Iannitelli et al., 2023).

The implication of these studies is that once neuromelanin accumulates past a tipping point, its beneficial activities cease and it results in a neuroinflammatory response that damages or kills cells. From this point of view, the association between cognition and neuromelanin intensity could simply reflect the fact that the tipping point has not yet been reached. The LC norepinephrine system is essential for maintaining optimal cognitive function with age. However, in our data, we found that older adults with below-average cognition did not have reduced neuromelanin, but lacked any relationship between neuromelanin intensity and cognition. This highlights that lower cognitive function has many sources beyond the LC, but maintaining higher cognitive function, especially related to fluid cognition, may depend on integrity of the LC norepinephrine system (Hayes & Petrov, 2016; Tsukahara et al., 2016; Tsukahara & Engle, 2021), though longitudinal data are needed to address this. This is an active topic of research, especially since the LC is the first structure to show tau pathology in the earliest stages of Alzheimer’s disease (Braak stage 0, (Braak et al., 2011), and could serve as either an early warning sign, or perhaps as a literal spreader of disease. However, the implications for LC damage go beyond AD, since substantial LC damage is also found in Parkinson’s disease (Zarow et al., 2003), and also other diseases such as multiple sclerosis (Polak et al., 2011), and pathological anxiety (Morris et al., 2020). What role neuromelanin, especially in its capacity to sequester environmental toxins, plays in all of these diseases is yet to be discovered.

Very recent work in the field is addressing crucial questions: whether individual neurons with the most neuromelanin are at the highest risk for imminent death (Nagatsu et al., 2023), whether neuromelanin content of an individual’s LC relates to its catecholamine production capacity and/or connectivity (Chen et al., 2023), and the true source of the neuromelanin MRI signal and how it can be processed more effectively to get a more precise look in vivo at the state of the LC norepinephrine system in humans (Trujillo et al., 2023). These studies should help to further clarify the meaning of neuromelanin as a functional biomarker and its ability to affect neural activity and health in the LC. Furthermore, other types of neuromelanin-sensitive MRI sequences, including magnetization transfer and quantitative susceptibility mapping sequences, are being tested and have the potential to bring converging (or diverging) evidence about the importance of neuromelanin. All of these advances should bring us closer to being able to understand when neuromelanin is beneficial vs. harmful and how the associated benefits or risks might be most productively managed.

## Methods

### Participants

We examined 96 participants between the ages of 19 and 86. For examination of neuromelanin across the lifespan, participants were broken up into 6 age groups: 18-29 (n = 32), 30-49 (n = 10), 50-59 (n = 9), 60-69 (n = 30), and 70+ (n = 15). Due to smaller sample size in middle-aged groups, for analyses by race and sex, participants were broken into 3 standard US Census groups instead: younger (19-44 years of age, n = 37, 27 female, 13 white, 5 Black), middle aged (45-64 years of age, n = 20, 12 female, 11 White, 6 Black), and older (65+ years of age, n = 37, 21 female, 31 White, 6 Black). Participants were screened for diagnosed cognitive impairment, neurological disease, head injury, ocular disease, and had vision and hearing that were normal or correctible to normal. All were fluent speakers of English. The sample included 55 white participants (57%), 16 Black participants (17%), 12 Asian participants (13%), 8 Latinx participants (8%), and 4 participants of other or multiple races or ethnicities (4%). The sample included 35 men (36%) and 61 women (64%). This research protocol was approved by the Cornell University Institutional Review Board for Human Participants with protocol number 1910009087.

### Cognitive assessment

Participants were assessed with the NIH Toolbox Cognition Battery (Hodes et al., 2013). Subsets used include Picture Sequence Memory Test, Flanker Inhibitory Control and Attention Test, Dimension Change Card Sort Test, List Sorting Working Memory Test, and Pattern Comparison Processing Speed Test. Uncorrected Standard Score was used to compare raw performance across age groups, and National Percentile (age-adjusted) was used to measure cognition relative to expectation. Participants were also assessed with the National Adult Reading Test (Venegas & Clark, 2011).

### MRI

Volunteers were scanned at the Cornell Magnetic Resonance Imaging Facility on a General Electric 3.0 Tesla MR750 scanner (Waukesha, WI, software version DV24 and DV29.1) with a 32 channel receive-only head coil. Magnitude images were saved to disk. Anatomical data were acquired with a T1-weighted MPRAGE sequence (TR = 7.7 ms; TE = 3.42 ms; 7° flip angle; 1.0 mm isotropic voxels, 176 slices). For the localization of the LC we used a protocol that provided the best LC contrast in a reasonably short scan time. It was inspired by previous publications with LC MRI protocols (Jacobs et al., 2021; Keren et al., 2009; Liu et al., 2020; Watanabe et al., 2019) and was optimized to fit our specific 3.0 Tesla MRI scanner model and its available pulse sequences. A 2D T1-weighted fast spin-echo sequence with ETL = 3 was used, with 18 slices of 3 mm thickness, field-of-view = 22 × 22 cm, acquisition and output matrix size = 512 × 512 (resolution = 0.43 × 0.43 mm), 60% FOV, TR = 701 ms, TE = 11.44 ms, refocusing flip angle = 120 degrees, 6 averages, bandwidth = 15.63 kHz, and extended dynamic range. Slices for this volume were oriented perpendicular to the brain stem to provide high resolution data in the axial plane, where the dimensions of the LC are smallest, and positioned to cover the most rostral portion of the pons. Participants laid supine on the scanner bed for all scans with their head supported and immobilized. Ear plugs, headphones, and a microphone were used to reduce scanner noise and to allow the participant to communicate with the experimenters.

### Analysis of images

In order to ensure that any voxels contained within the 4th ventricle were not used in neuromelanin analysis, MPRAGE images were processed using FreeSurfer *recon-all* (Fischl, 2012) in order to produce an automated ROI corresponding to the 4th ventricle. The MPRAGE image, and corresponding parcellation, was then aligned with the TSE image with AFNI (Cox, 1996) using the *3dAllineate* function. After definition of the LC ROI, any voxels overlapping with the 4th ventricle were removed. Measurement of neuromelanin intensity for each participant was then conducted as described below using the TSE image.

### Probabilistic map

A probabilistic map illustrating the location and distribution of the LC ROIs in all participants was created by registering each person’s LC ROI to MNI152 space. To do so, alignment parameters were calculated for each individual by combining the inverse of the transformation that aligned the native MPRAGE and TSE images and the nonlinear transformation from the individual’s native MPRAGE to MNI152 standard calculated by AFNI functions *3dQwarp* and *3dNwarpApply*. Successful alignment of the brainstem to the individual MPRAGE and the MNI152 template was visually confirmed. LC ROIs in MNI152 space were thresholded to remove scattered non-zero voxels introduced by the nonlinear warping process. A heatmap was then created using AFNI function *3dMerge*.

### Measurement of NM intensity

NM intensity was measured by exactly following the procedure described in our previous published work (Turker et al., 2021). Briefly, raw data was processed for correction of artifacts and optimized for contrast. Then, LC ROIs were hand-drawn using *FreeView* software according to clear guidelines (Turker et al., 2021) by two independent raters. Only the overlap of both ROIs was used, less any voxels belonging to the 4th ventricle mask, after ensuring that the Dice similarity coefficient between raters was greater than 0.6. If the coefficient was less than 0.6, both raters reviewed the ROI drawing procedure and independently redid the ROI. This was necessary for only 4 out of 96 participants. The average Dice coefficient was 0.84 for all participants, 0.839 for participants under age 60 and 0.841 for participants over 60, demonstrating good agreement of the ROI drawing procedure across age groups.

After creation of the ROIs, neuromelanin images were then rescaled to a range of 6 standard deviations. This transformation was chosen to be as similar as possible to min-max scaling (min 0, max 1), without undue influence of extreme outliers caused by scanner noise. The scaling range was calculated using all voxels in the brainstem (as opposed to a specific reference region of the pontine tegmentum) in slices where the LC was defined, except for those within the LC ROI. The transformation was then applied to all voxels, including those within the LC, such that the neuromelanin intensity could then be interpreted relative to the range of non-LC values within all slices in which LC was observed. For example, a value of 2 indicates that the LC was twice as intense as the maximum (actually 3 SD above mean) of the non-LC values. Neuromelanin intensity was defined as the sum of all voxel values within the LC mask divided by the number of voxels. Rostral vs. caudal regions were defined by a 50/50 split along the rostral-caudal axis of the overall length of the individually defined LC ROI for each person. There was one participant with an outlier value, an average neuromelanin intensity 5.8 SD above the total sample average. This participant was removed from analysis, but inclusion or exclusion of this individual did not change the result of any statistical tests.

## Statistics

Linear models were run in R using the *lme4* package (Bates et al., 2015). Summary statistics such as p values and F statistics, were calculated using the *anova* function, while p values and t statistics for levels within a predictor (e.g., to determine which age groups differ from the reference group in a model in which age group was a significant predictor) were calculated using the *summary* function. One general additive model was produced using the *mgcv* package (Sørensen et al., 2021).

## Acknowledgements

The authors wish to thank Henning Voss of the Cornell MRI Facility for consultation, optimization and troubleshooting in the use of neuromelanin-sensitive MRI sequences.

## Data Availability Statement

For privacy reasons, neither full anatomical MRI scans nor full neuromelanin-sensitive MRI scans can be made available. However, neuromelanin-sensitive MRI scans limited to the brainstem, in addition to all cognitive and demographic data necessary to replicate the analyses described here, are available at the Open Science Framework at https://osf.io/tf9br

